# Chronic CNS Pathology is Associated with Abnormal Collagen Deposition and Fibrotic-like Changes

**DOI:** 10.1101/2021.07.06.451298

**Authors:** Daphne Palacios Macapagal, Jennifer Cann, Georgia Cresswell, Kamelia Zerrouki, Karma Dacosta, Jingya Wang, Jane Connor, Todd S. Davidson

## Abstract

Multiple sclerosis is a chronic debilitating disease of the CNS. The relapsing remitting form of the disease is driven by CNS directed inflammation. However, in the progressive forms of the disease, inflammation has abated and the underlying pathology is less well understood. In this paper, we show that chronic lesions in progressive MS are associated with fibrotic changes, a type of pathology that has previously not thought to occur in the CNS. In an animal model of chronic MS, late stage disease contains no inflammatory infiltrates and is instead characterized by collagen deposition that is histologically similar to fibrosis. In human MS samples, chronic, but not acute lesions, are devoid of inflammatory infiltrates and instead contain significant collagen deposition. Furthermore, we demonstrate that both mouse and human astrocytes are the cellular source of collagen. These results suggest that anti-fibrotic therapy may be beneficial in the treatment of progressive MS.

## Introduction

Multiple sclerosis is an autoimmune disease of the central nervous system. Approximately 80-85% of patients with MS present with a relapsing-remitting (RRMS) course of disease characterized by episodes of disability followed by a period of remission.[1] After a period of years, RRMS can convert into a more progressive disease (secondary progressive, SPMS) in which the patients steadily decline. The remaining 15-20% of patients with MS will present with a progressive disease from the onset (primary progressive, PPMS)[1, 2] RRMS is primarily an inflammatory disease wherein T and B cells, along with macrophages infiltrate the central nervous syste (CNS) and cause tissue damage. Such lesions can readily be detected by MRI as gadolinium (Gd) enhancing lesions. Though rare, these lesions are not observed in the progressive forms of disease[3]. In addition, RRMS is amenable to treatment with anti-inflammatory treatments, whereas such treatments are generally considered ineffective in treating SPMS and PPMS.

Most MS research has been focused on RRMS as it is the most common form of the disease. Less is known about the progressive forms. One hurdle in studying progressive MS has been the lack of appropriate in vivo models. Experimental autoimmune encephalomyelitis (EAE) has been widely used as an animal model of MS. EAE is induced in susceptible strains of mice by immunization with various myelin derived peptides in adjuvant, resulting in T cell activation, migration into the spinal cord, and subsequent paralysis. This approach has been useful in modeling the immune driven aspects of RRMS, but its highly inflammatory nature would seem to make it inappropriate as a model of progressive MS.

There does not appear to be major genetic differences between RRMS and progressive MS in terms of susceptibility loci, etc. This suggests that the various forms of disease represent stages of the same disease, as opposed to distinct disease entities. We reasoned that EAE may have similar, unappreciated aspects. For example, immunization of B6 mice with MOG35-55 in CFA induces a chronic disease driven by CD4+ T cells. However, the vast amount of research on this model has only focused on the acute phase of disease, while the chronic phase has been relatively ignored.

Here, we take an unbiased gene expression approach to characterizing the chronic phase of EAE in the B6 mouse to determine its suitability as a model of SPMS. We determined that chronic EAE is not driven by inflammation, but instead is driven primarily by abnormal remodeling of the extracellular matrix reminiscent of fibrosis. We confirm these findings in human samples and demonstrate their association with various forms of MS. This work demonstrates a previously unappreciated aspect of progressive MS and suggests avenues to future therapies.

## Results

Experimental autoimmune encephalomyelitis in the C57Bl/6 mouse is a well characterized model of autoimmune mediated neuroinflammation. Often used to demonstrate aspects of multiple sclerosis, it has been studied at the genetic, cellular and molecular level where it has been shown to be driven by autoreactive CD4+ T cells and inflammatory macrophages. Recent advances in genomics technology has enabled characterization of gene expression profiles at unprecedented scale and resolution. Most gene expression studies of EAE have focused on highly purified populations of cells, or even single cell analysis. Moreover, almost all studies have focused on early time points (days 11-23). While analysis of purified cell populations or single cell analysis allows deep understanding of the function of a particular cell type, it necessarily omits changes occurring at the global level.

We sought to identify novel pathways regulated during the course of EAE. B6 mice were immunized with MOG35-55 and spinal cords were collected at various time points (Figure 1a). We grouped the disease stage into three phases, pre-disease (days 0 and 7), acute disease (days 16 and 23), and chronic disease (days 31 and later). As we were attempting to identify novel pathways associated with disease progression, and not simply changes in gene expression levels due to differences in severity, we carefully selected mice with a clinical score of 3, characterized by limp tail and paralysis of the hinds limbs, on a scale of 0 to 5 (0=normal; 5=moribund) in the acute and chronic phases of disease.

**Figure 1:**
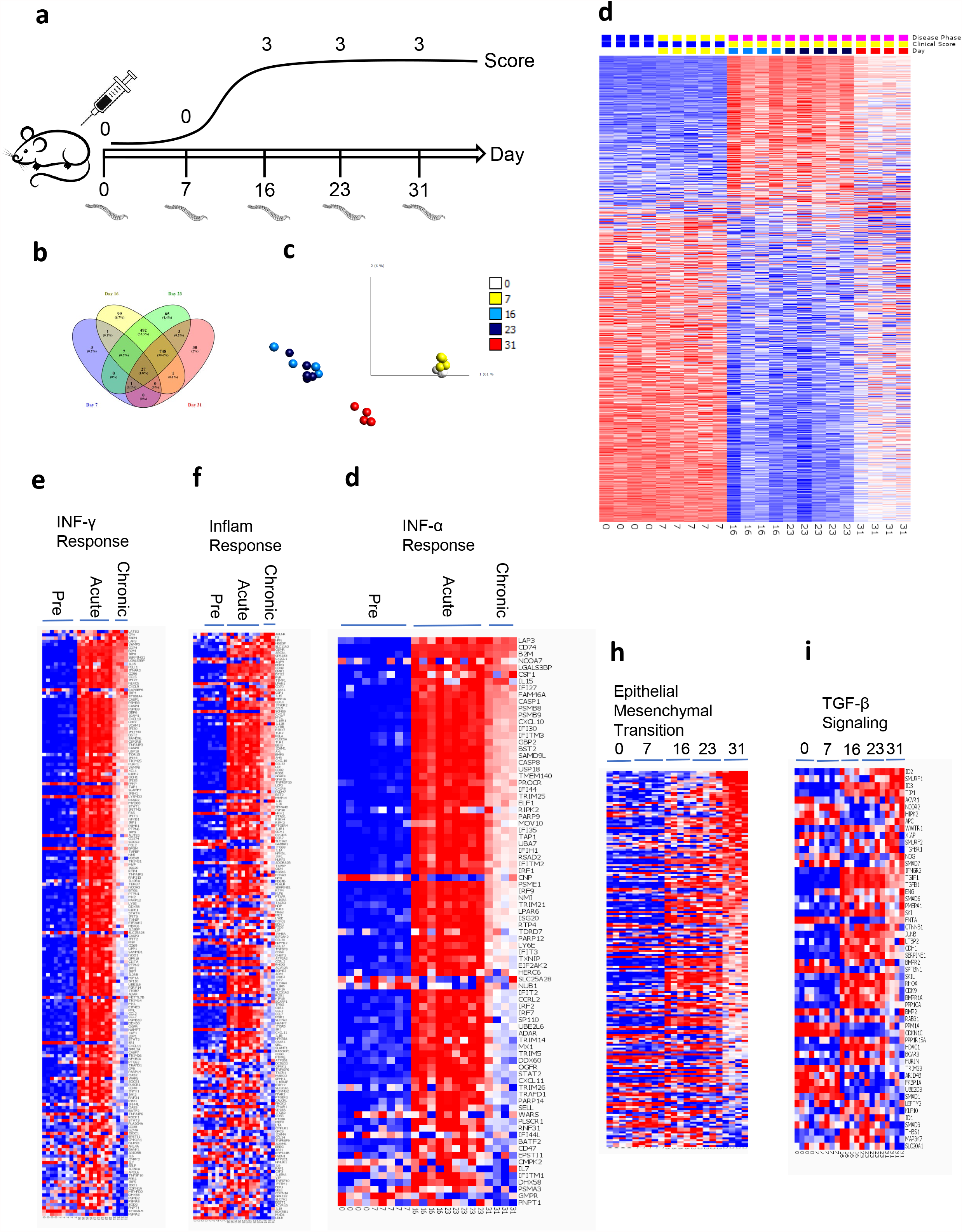
A, Schematic of experimental design. B6 mice were immunized with MOG35-55 on day zero and spinal cords were harvested on the indicated days. Mice sacrificed on days 0 and 7 had a clinical score of 0, while only mice with a clinical score of 3 were selected at the following time points. B, Venn diagram showing the overlap of regulated genes at the indicated time points. C, PCA plot showing clustering of individual mice at the indicated time points for genes regulated at FDR < 0.05. D. Heatmap display of using the same parameters as in C. E-I, Heatmaps of genes regulated in GSEA genes sets according to disease phase.

RNA from the spinal cords of individual mice was analyzed by gene array. The resulting data were filtered to genes with expression levels changed greater than 1.5 fold (normalized to 18s) and with p values less than 0.05. This resulted in a list of 1477 genes that were significantly regulated. The overlap of genes amongst the various time points can be seen in Figure 1b, with the plurality of genes located in the overlap of days 16, 23, and 31 and thus representing genes that distinguish inflamed from non-inflamed CNS. PCA analysis of the significantly regulated genes identified three discreet populations (Figure 1c). Pre-disease spinal cords clustered tightly together, suggesting minimal to no global changes at the gene expression level preceding clinically evident disease. Spinal cords from the acute phase of disease also clustered tightly together, as was expected considering we selected mice with identical clinical scores. Gene expression data suggest that spinal cords from the chronic time point clustered separately from either of the aforementioned populations. This was somewhat unexpected, as we selected mice with identical clinical scores to those in the acute phase of disease. This indicates that although mice in the acute and chronic phase of disease have identical clinical presentations, the underlying gene expression patterns are dramatically different.

This can be visually seen in Figure 1d, where the gene expression patterns between pre-disease and acute disease exhibit an almost inverse relationship. This was likely due to the influx of inflammatory cells that are not normally present in the CNS. The expression pattern at the chronic phase of the disease shows a downregulation of certain genes that were later upregulated in the acute phase. Likewise, an inverse upregulation of a certain number of genes that were downregulated during the acute phase.

We next used Gene Set Enrichment Analysis (GSEA) to identify pathways regulated during different phases of disease. As expected, a number of pathways involved in the inflammatory response were upregulated in acute disease relative to pre-disease. These pathways included the IFN-γ response (Figure 1e) and the inflammatory response gene set (Figure 1f). Less well studied in EAE has been IFN-α, but this gene set was also clearly regulated (Figure 1g). However, closer inspection of this particular gene set shows significant overlap with the genes regulated in the IFN-γ gene set, suggesting that IFN-α may not be the driving factor. Importantly, these three gene sets showed upregulation of key inflammatory genes during the acute disease stage that were subsequently down regulated in the chronic phase.

To this point we had identified a clear difference at the gene expression level between acute and chronic disease. The majority of the regulated genes were associated with inflammation and were downregulated at the chronic phase of the disease. Since mice in the acute and chronic phases had identical clinical scores, it was unclear what pathways were being activated to drive pathology at the later stage. We therefore identified GSEA pathways which were upregulated in the chronic phase, but not during the acute or pre-disease stages. We identified only one such GSEA gene set which clearly fit this criteria, epithelial mesenchymal transition (Figure 1h). EMT (Epithelial Mesenchymal Transition) is a pathway in which epithelial cells transform into mesenchymal cells and is one of several mechanisms which can lead to the development of fibrosis. A key driver of EMT is TGF-β signaling. We identified upstream mediators of TGF-β signaling in acute disease and downstream mediators of TGF-β signaling in chronic disease which clearly fits this criteria (Figure 1i).

We first sought to confirm our gene array data and examined more closely the expression profiles of fibrosis associated genes in the CNS. To this end, we designed a Fluidigm panel with an array of fibrosis related probes, including multiple collagen isoforms, genes involved in collagen organization, as well as a number of cytokines known from other organ systems to be involved in inducing collagen synthesis. The panel also included a number of neuronal specific markers and genes involved in the inflammatory response. This analysis included RNA from the original samples used to run the gene array, as well as additional samples isolated from an independent experiment.

Hierarchical clustering of the Fluidigm results (Figure 2a) demonstrated clusters of genes regulated at various stages of disease. We confirmed upregulation of inflammatory genes in acute disease relative to pre-disease, followed by downregulation in chronic disease. We also identified a cluster of neuronal markers that were down regulated in acute disease relative to pre-disease. This could be due to death of neurons or could be reflective of the fact that the majority of the RNA in these samples were derived from inflammatory cells thus diluting the neuronal cells RNA in the sample as a whole. Importantly, we saw upregulation of a number of collagen genes, including Col1a1, Col1a2, Col3a1 and Col4a1, all of which are important constituents of fibrotic lesions (Figure 2b). We also saw upregulation of a number of genes involved in organization of the collagen matrix, including Lumican, FAP, FMOD and Serpin1. TGF-β and its receptors TGFβR1 and TGFβR2 were upregulated in both acute and chronic disease. IL-13, also known to be involved in the progression of fibrosis, was upregulated in acute disease along with its receptor IL-13Ra1, while its decoy receptor IL-13Ra2 was downregulated.

**Figure 2:**
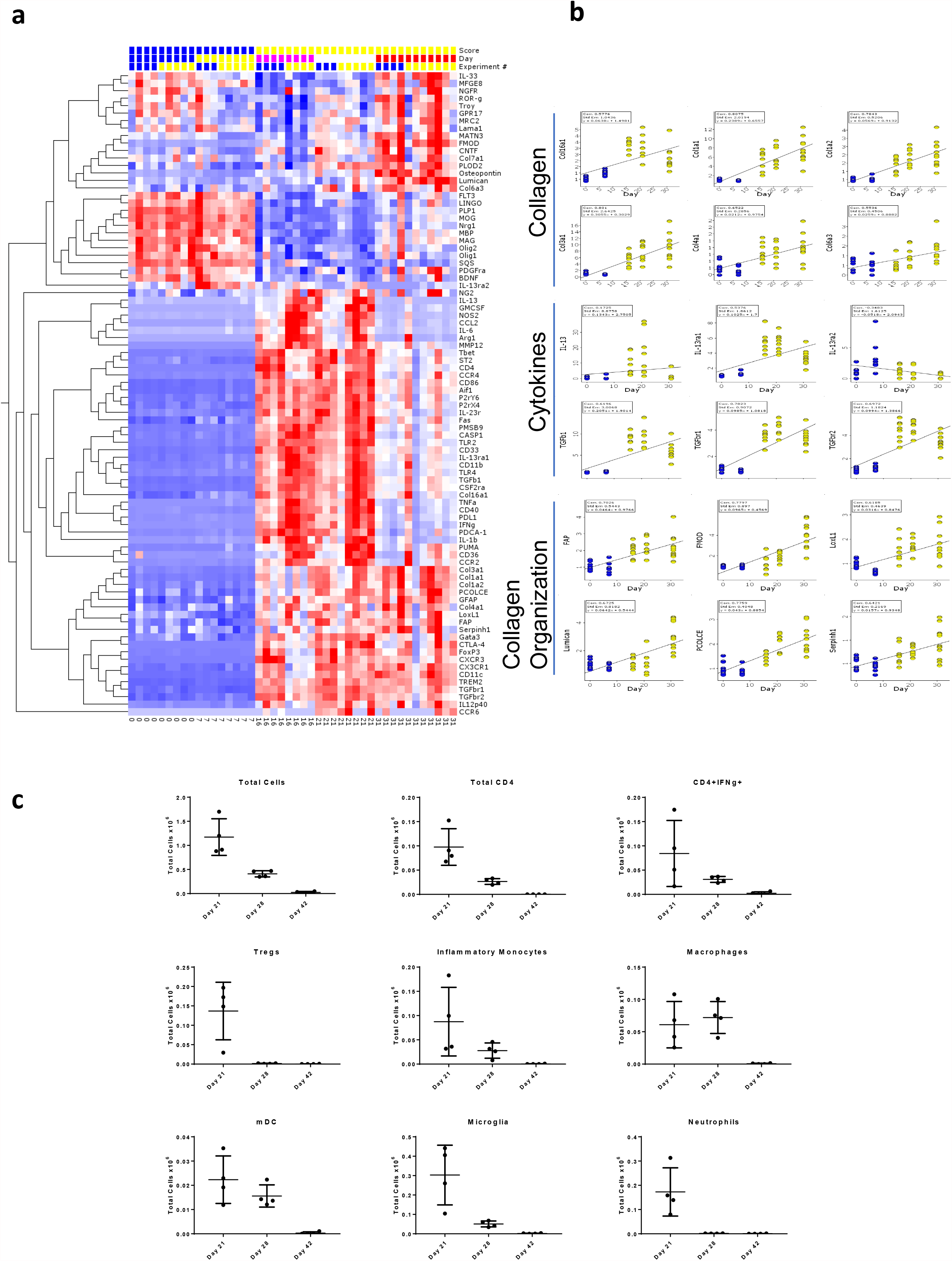
A, Fluidigm analysis of genes identified in the gene array. Genes were selected based on their appearance in the GSEA gene sets related to collagen, cytokines, and collagen organization. B, Kinetic profiles of select gene expression levels as determined by Fluidigm analysis. Pre-disease indicated in blue, active disease indicated in yellow. C, Total cell counts of the indicated populations in the spinal cord as determined by flow cytometry. Total cells are all CD45+ cells, total CD4 is CD45+CD4+, CD4+IFNg+ is CD45+CD4+IFNg+, Tregs are CD45+CD4+Foxp3+, Inflammatory monocytes are CD45+ CD11b^+^ CD11c^-^ Ly6G^-^ Ly6C^hi^, macrophages are CD45+ CD11b^+^ CD11c^-^ Ly6G^-^.

EAE is known to be driven by CD4+ Th1 cells and inflammatory macrophages. Our gene expression data suggests that inflammation has abated during the chronic stage of the disease. We confirmed this by flow cytometric analysis of CNS infiltrating leukocytes at various time points (Figure 2c). As stated before, we only selected mice with a clinical score of 3. We found large numbers of CD45+ cells in the CNS of mice on day 21, the peak of EAE disease. This number declined by day 28, and we found almost no CD45+ cells in the CNS on day 42, despite the fact that the mice still had significant clinical disease. Subset analysis of CD45+ cells identified a similar pattern for CD4+IFN-γ+ cells, as well as inflammatory monocytes, macrophages, and mDCs.

Our current understanding of EAE is that it is driven almost exclusively by inflammation. However, these data suggest that during the chronic stage of the disease inflammation has abated and fibrosis drives pathology. This is a common pathway in many diseases, but to our knowledge fibrosis has never been documented in the CNS. We therefore sought to identify fibrotic lesions in the spinal cords of chronic stage EAE mice at the histological level. We isolated spinal cords from mice with EAE at various time points and stained them with Sirius Red (Figure 3a), Masson’s Trichrome (Figure 3b) and H&E (Figure 3c). Sirius red and Masson’s trichrome stained sections showed no evidence of fibrosis or abnormal collagen deposition up to day 23. By day 31, abnormal accumulations of collagen were present along the meningeal interface and by day 41 abnormal collagen deposition was clearly present throughout the parenchyma. The pattern of collagen deposition was clearly abnormal for the CNS, but it was not a classic fibrosis morphology with compact bundles of mature collagen. Perivascular collagen deposition was clearly evident, but the majority of collagen was in the parenchyma and had a fibrillar deposition. H&E sections demonstrated inflammatory infiltrates in acute disease that had abated by the chronic phase.

**Figure 3:**
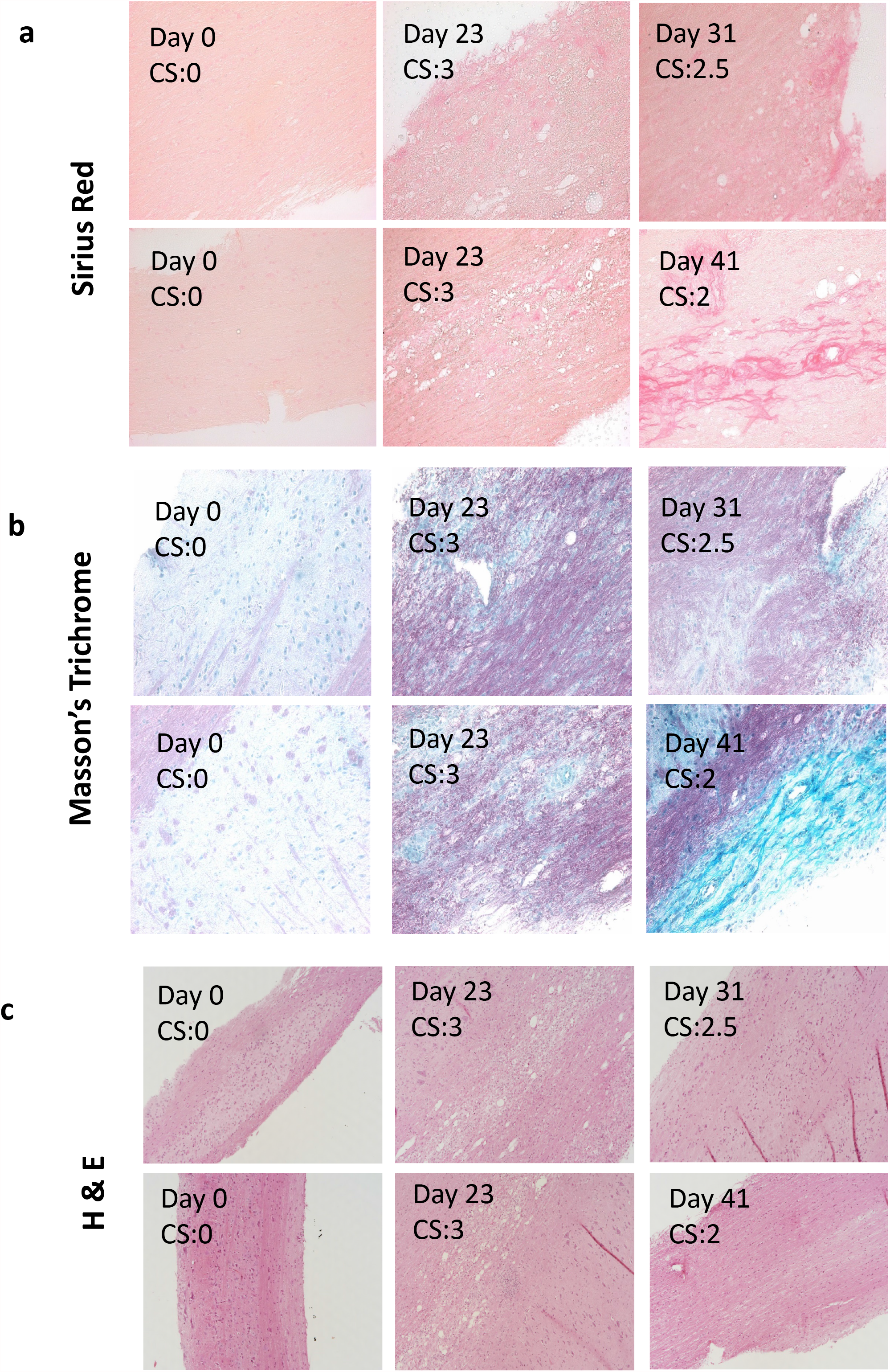
B6 mice were immunized for EAE and sacrificed on the indicated days. The clinical score (CS) of each individual animal is indicated. Serial sections were stained with Sirius Red, A, Masson’s Trichrome, B, or H&E, C, to identify areas of fibrosis and to characterize histologically.

In most organ systems, fibrosis is mediated by activated fibroblasts. However, the CNS is devoid of such cells, raising the question of which cells produce collagen. We approached this question by co-staining for collagen and various cell types in the CNS of mice with EAE. We found by brightfield microscopy that collagen was associated with astrocytes in diseased, but not normal tissues (Figure 4a). In normal tissue, collagen was associated solely with vessels, and astrocytes were distributed evenly throughout the tissue with characteristic stellate morphology. In mice with chronic EAE, the astrocytes significantly changed their morphology displaying much smaller cell bodies with a fibrillar morphology. To more conclusively demonstrate colocalization of collagen and astrocytes, we analyzed the sections by confocal microscopy (Figure 4b). Non-vascular collagen was found to be significantly, although not exclusively associated with GFAP+ cells.

**Figure 4:**
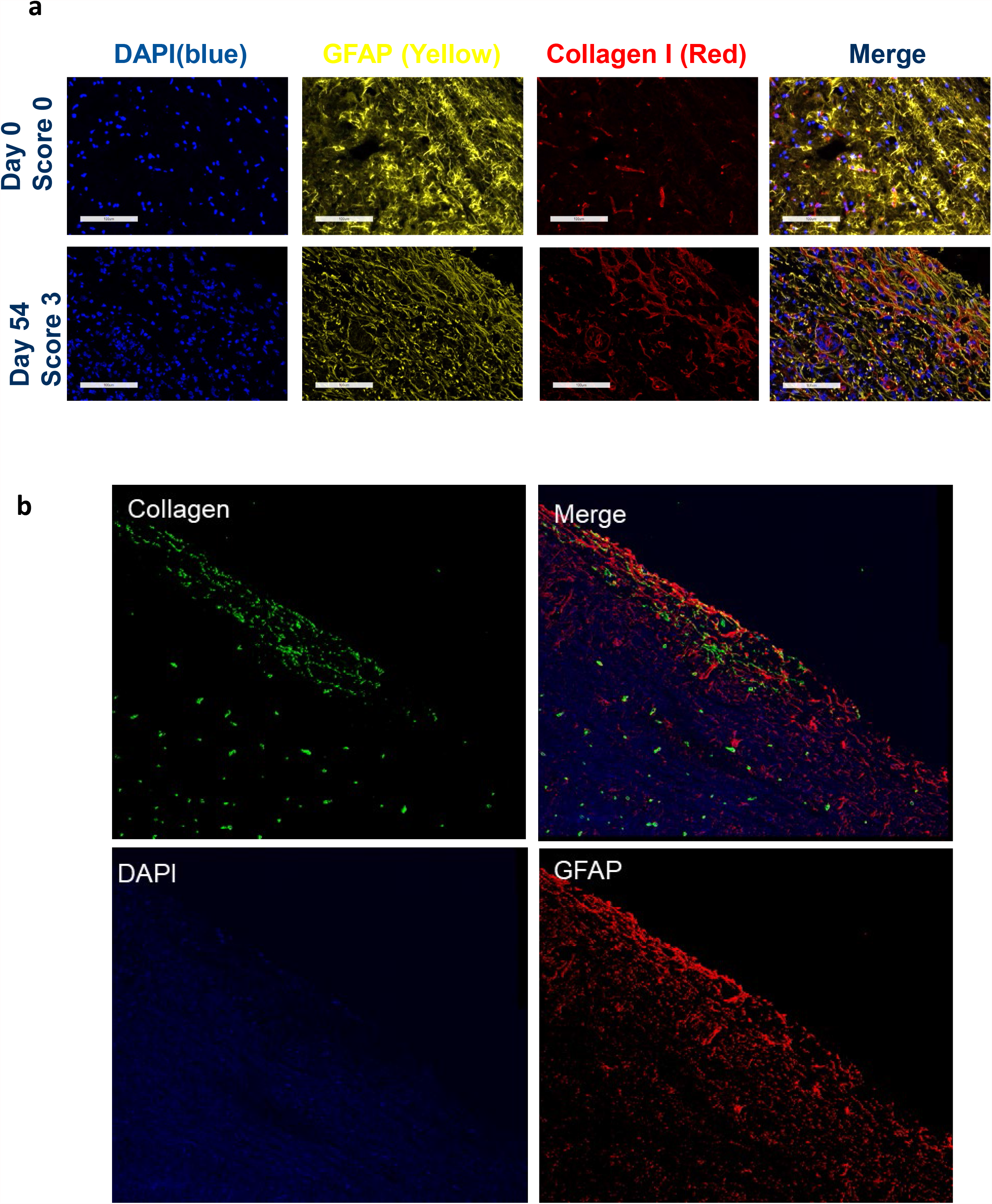
Fluorescent microscopy demonstrates co-localization of collagen to GFAP+ astrocytes. B6 mice were immunized for EAE and sacrificed at either day 0 (clinical score of 0) or day 54 (clinical score of 3). Spinal cords were collected and processed for fluorescent microscopy by staining with GFAP (yellow), Collagen 1 (red) and DAPI (blue) and analyzed by fluorescent microscopy, A, or by confocal microscopy, B.

EAE in the B6 mouse is a chronic disease. We have thus far shown that in the later stages of disease, inflammation has abated and the prevailing pathology is fibrotic. We next asked to what extent these fibrotic changes are responsible for the clinical pathology. We and others have previously shown that neutralizing GM-CSF reverses established disease [4] [5] and [6]. Mice treated with anti-GMCSFR have reduced T cell and inflammatory macrophage infiltrates [4]. We treated mice with established EAE with CAM3003 (anti-GMCSFR) or isotype control and allowed the mice to recover. Spinal cords from mice treated with CAM3003 had fewer Sirius red lesions than mice treated with isotype control (Figure 5a and b). These results suggest that in the chronic stage of disease, GM-CSF signaling is required for the establishment of fibrosis and not the progression of inflammation.

**Figure 5:**
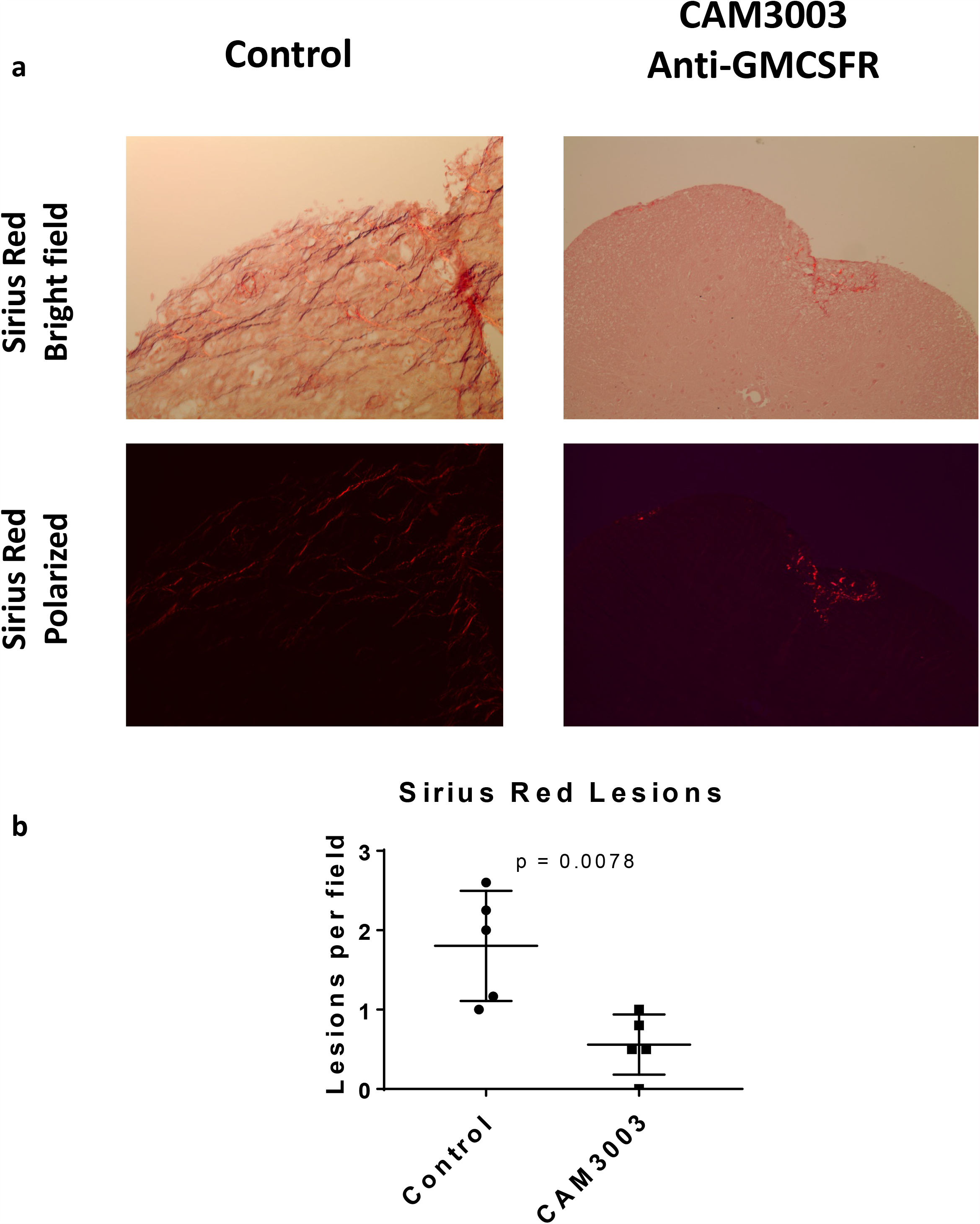
Anti-GMCSFR prevents the establishment of fibrosis in the CNS. B6 mice were immunized for EAE. At peak of disease (day 14) mice were treated intraperitoneally every other day with 10mg/kg anti-GMCSFR (CAM3003) or control until disease subsided. Spinal cords were collected, paraffin embedded and stained with Sirius Red to detect collagen deposition. A, representative images obtained under either brightfield or polarized conditions at 20X magnification. B, each dot represents the average lesions per field for an individual mouse obtained under 10X magnification for mice that received either CAM3003 or control treatments.

To assess collagen content in MS, we immunohistochemically stained sections of brain from human MS patients for collagen-1a (Figure 6a). In all areas of the brain, strong reactivity for collagen-1a was seen in the basement membranes of blood vessels, including small capillaries throughout the neuropil. In chronic demyelinated and sclerotic foci within the white matter, additional fine fibrillar structures and mononuclear cells with fusiform or stellate morphology were also strongly positive for collagen-1a. To determine if these might represent reactive endothelium or attempts at neovascularization, we immunohistochemically stained a separate nearby section for the endothelial cell marker, CD31 (Figure 6b) [7, 8]. In all areas of the brain, moderate to strong reactivity for CD31 was seen in endothelial cells, colocalizing with the underlying collagen-1a+ basement membranes. However, there was no CD31 staining of the fine fibrillar structures or mononuclear cells with fusiform morphology within the sclerotic foci that were strongly positive for collagen-1a. To determine if these might represent astrocytes cell bodies and their processes, we immunohistochemically stained a nearby section with GFAP (Figure 6c). In non-sclerotic areas of the brain, astrocyte cell bodies and their processes were strongly positive for GFAP, but the surrounding cells and neuropil were negative. In the sclerotic foci, strongly positive astrocyte cell bodies and their processes were densely packed and colocalized with the collagen-1a+ fine fibrillar structures and mononuclear cells. This suggests that within the glial scar, some astrocytes are actively producing and depositing collagen-1a.

**Figure 6:**
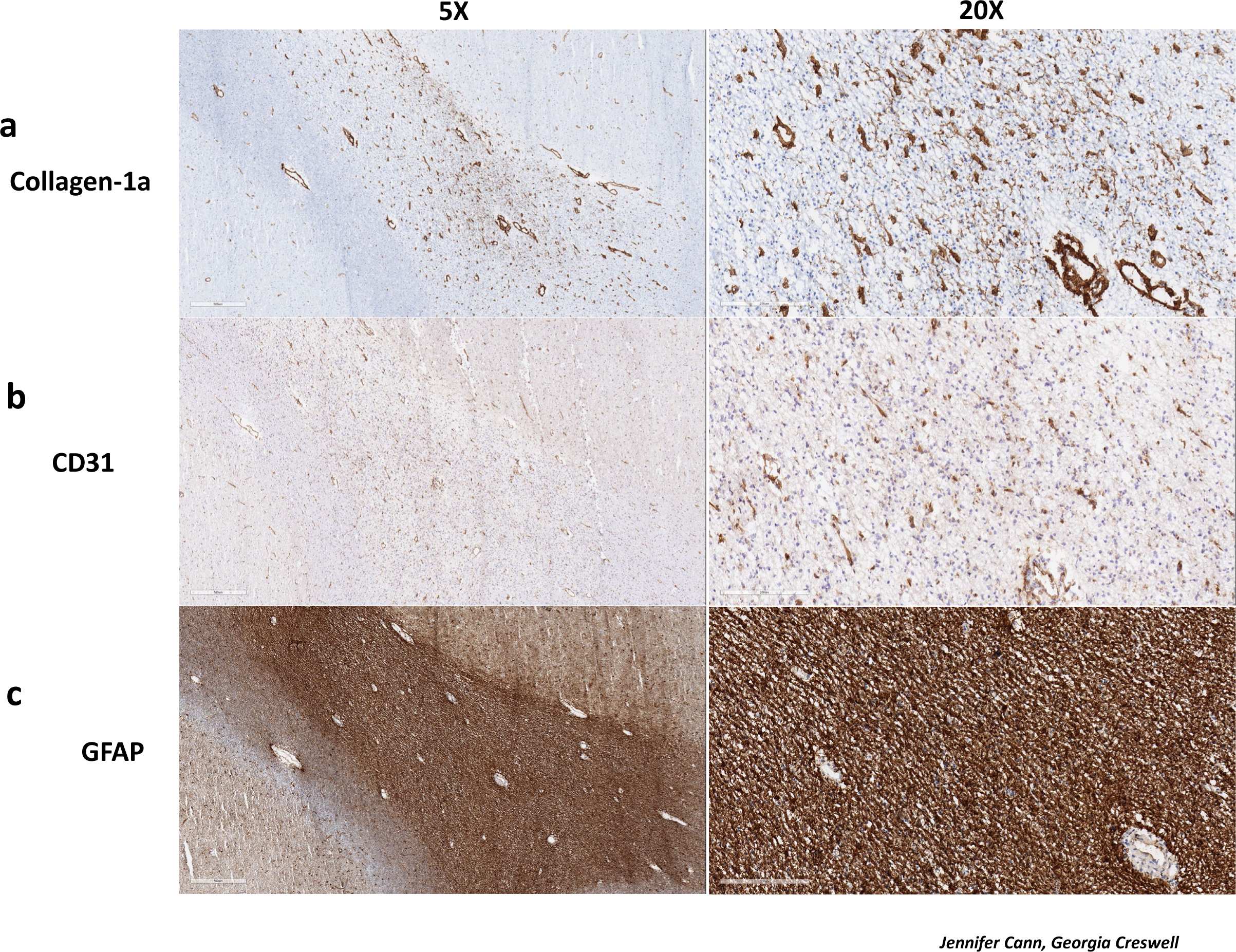
Collagen deposition in MS plaques. Samples of human MS tissue were stained for collagen-1a, A, CD31, B, or GFAP, C and evaluated by brightfield microscopy.

## Discussion

In this study we identified a strong fibrosis associated gene signature in late stage EAE. We demonstrate abnormal collagen deposition at the histological level and show that similar pathological changes happen in human MS samples. These changes only occurred at very late time points in the course of the disease and coincide with a marked reduction in inflammatory infiltrates. Therapeutic intervention with GM-CSFR blocking antibodies ameliorated inflammation and prevented late stage fibrotic changes. Finally, we demonstrated that astrocytes are the source of the abnormal collagen deposition.

Fibrosis is the formation of fibrous connective tissue as the result of deposition of extracellular matrix proteins [9, 10]. Fibrosis has been demonstrated in the meninges [11] [12] yet is thought to not occur in the CNS. Indeed, we found a very distinct pattern of collagen deposition in CNS lesions. These lesions were highly fibrillar and located in chronic resolving lesions. We had considered that the collagen deposition we were seeing was related to neovascularization as such processes are known to take place in a number of other pathological process in the CNS such as traumatic brain injury, stroke and ocular neovascularization [13] [14]. However, we found no co-localization of CD31 with collagen producing cells in MS lesions, suggesting that the collagen deposition observed was not related to neovascularization.

The deposition of collagen could negatively influence reparative processes. For example, oligodendrocyte precursor cells are unable to differentiate into mature myelinating oligodendrocytes on stiff matrices [15]. The role of matrix stiffness has been investigated much more extensively in various fibrotic diseases such as those of the lung and skin [16-19]. In these instances, stiff collagen matrices dramatically alter the phenotype of resident stromal cells. Our results would suggest that abnormal collagen deposition in the CNS impairs normal physiology.

Scarring is a well appreciated feature in MS plaques, but to our knowledge this study is the first demonstration that MS plaques are associated with abnormal collagen deposition. Scarring is associated with astrocytosis and recently astrocytes were shown to adopt a neurotoxic phenotype – termed A1 astrocytes – in MS lesions [20]. Itoh et al have shown in late stage EAE that astrocytes downregulate genes involved in the cholesterol biosynthesis pathway and upregulate proinflammatory genes [21]. In addition, numerous studies have demonstrated dysfunctional astrocytes associated with axonal degeneration [22]. These studies demonstrate that astrocytes play a detrimental role in chronic CNS disease. Our work expands on this concept by showing that astrocytes produce collagen and that this collagen prevents productive repair.

In this study we investigated gene expression changes in the spinal cord of mice during different stages of EAE. We carefully selected mice with identical disease scores so that any changes we uncovered were due to the stage of the disease, and not related to severity. We chose to analyze whole spinal cord as opposed to isolated cell populations. Such an approach limits the ability to ascribe any gene changes to a specific cell type but allowed us to identify pathways occurring at a more global scale that may have been missed had we chosen to focus on individual cell populations. GSEA analysis identified pathways that were differentially regulated at various stages of the disease. As expected, acute and peak disease states were associated with a massive upregulation of proinflammatory genes. EAE is well known to be driven by inflammation, yet at the chronic stage of the disease inflammation has completely abated and despite the fact that the mice continued to display clinical symptoms. The main pathways that were upregulated at these later time points were associated with fibrosis, suggesting that this is the predominant pathway associated with pathology at this stage.

Relapsing remitting MS is highly inflammatory and immunomodulatory treatments are relatively successful at controlling disease [23, 24]. The progressive forms of MS are less inflammatory and have proven to be relatively refractory to immunomodulatory interventions [1, 25]. Progressive MS lesions are characterized by neurodegenerative changes including axonal damage and gliosis [1]. Indeed, it was in these lesions that we identified high levels of abnormal fibrotic changes. Although there are currently no truly effective treatments for progressive MS, a considerable amount of research has been focused on neuroregenerative approaches. Our data suggest that antifibrotic approaches may be beneficial as well. Indeed, it may be that antifibrotic interventions will be required for successful neuroregeneration to occur.

## Methods

### Induction of Active EAE Disease and Scoring

EAE was induced in 6-9 week old female C57BL/6 mice. On day 0, animals were immunized subcutaneously in the flanks with 400µg of myelin oligodendrocyte glycoprotein 35-55 (MOG 33-55) (MEVGWYRSPFSRVVHLYRNGK; Anaspec Inc. Catalog No. AS-60130-1) in a 200ul emulsion of Complete Freund’s Adjuvant (4mg/ml Mycobacterium tuberculosis; Difco Labs Catalog No. 263810). On day 0 and day 2, 350ng of Pertussis toxin (Bordatella pertussis; Calbiochem Catalog No 516 560-50ug) was injected intra-peritoneally (i.p.). EAE scores were assessed daily for clinical signs of EAE in a blinded fashion. Animals were scored as follows: 0=normal; 1=limp tail; 2=hind leg paralysis of one leg or difficulty walking/ataxia; 3=paralysis of both hindlimbs (paraparalysis) and tail; 4=hindlimbs paraparalysis and one forelimb weakness; 5=moribund (requires sacrifice). All animal procedures were approved by the IACUC board (Institutional Animal Care and Use Committee) of Medimmune Inc. and the protocols followed were in accordance to the guidelines with the Animal Welfare Act (AWA).

### CNS tissue isolation

Animals were asphyxiated using a lethal dose of CO2. Animals showing no signs of pedal and palpebral reflex were first perfused intracardially using room temperature saline. The spinal cords were isolated and frozen on dry ice. Spinal cords were collected at naïve (score 0, day 0), onset (score 0, day 7), peak (score 3-4, day 16), post peak (score 3, day 23) and chronic (score 3 or higher, day 30-onward) clinical points. These samples were stored at -80°C until further processing for RNA extraction. Samples that were to be processed for immunohistochemistry, Sirius red or Masson’s Trichrome were placed in 10% formalin for 24 hours followed by paraffin embedding. Other spinal cords were isolated and frozen at -80°C in Tissue-Tek ^®^ O.C.T. compound. These samples will be further processed for IHC.

### RNA extraction from frozen spinal cords

Each frozen spinal cord from individual mice was processed separately for RNA extraction. Frozen material was placed into a 2mL RNASE/DNASE free Lysing Matrix D tube filled with beads (MP Biomedicals Catalog No. 6913-500) for disruption. Instructions were followed using the Qiagen Rneasy^®^ Lipid Tissue Mini Kit (Qiagen Catalog No. 74804) thereafter. Briefly, the spinal cords were homogenized using 1ml Qiazol^®^ Lysis Reagent (Qiagen Catalog No.79306) and FastPrep-24 5G Homogenizer (MP Biomedicals) for 40 seconds. The lysate was incubated at room temperature for 5 min, topped off with 200 µl of chloroform (Sigma-Aldrich Catalog No.C2432-500mL), shaken, incubated for 2-3min at RT, and centrifuged (12,000g at 4°C for 15min). The upper phase was transferred to 1 volume of 70% ethanol and placed into an RNeasy Mini spin column for centrifugation (8,000g at RT for 15 sec). The flow through was discarded and an optional on-column DNAse digestion protocol was followed. Briefly, RNeasy^®^ Mini spin column was washed with 350 µl of RW1 buffer then centrifuged (8,000g at RT for 15 sec). Then 80µl of DNase I incubation mix was added to the spin column (15-25°C for 15 min) and a final wash of 350 µl of RW1 buffer followed with centrifugation (8,000g at RT for 15 sec). RNA was extracted following the final steps of the Qiagen RNeasy^®^ Lipid Tissue Mini Kit as per manufacturer’s instructions. RNA samples were then prepared for Fluidigm^®^ Biomark HD array preparation.

### Fluidigm Biomark HD array preparation

Total extracted RNA was isolated from each mice spinal cord and used in preparation for Fluidigm^®^ Biomark array. The preparation of RNA was performed with the following steps: (1) cDNA synthesis; (2) cDNA preamp PCR; (3) Biomark HD priming; (4) Biomark assay loading. For cDNA synthesis, 50 ng of total RNA from each sample (3 mice per group) was used to initiate the PCR using SuperScript III Reverse Transcriptase (Invitrogen Catalog No. 18080-093). cDNA synthesis was performed as per manufacturer’s instructions. Preamplification of cDNA followed using the newly generated cDNA and TaqMan^®^ PreAmp Master Mix Kit (Applied Biosystem Catalog No. 4384556). The manufacturer’s suggested protocol was followed with minor modifications. 20X TaqMan^®^ Gene Expression assays (Invitrogen) were combine to a final .2X concentration. These will be used as a pooled assay mix for the PCR. Combined with the TaqMan^®^ PreAmp Master Mix and newly generated cDNA, the PCR reaction follows a hold of 95°C for 10 min and a 14X cycled reaction of 95°C for 15 sec and 60°C for 4 min. The final solution is dilute 1:5 with DNA Suspension Buffer (10mM Tris, .1mM EDTA, ph8.0, sterile and DNase/Rnase Tested) (Teknova, Catalog No. T0220).

After samples were preamped, the cDNA will be loaded onto a primed Fluidigm 96.96 Dynamic Array IFC (integrated fluid circuit). Manufacturer’s instructions were followed. Briefly, the 96.96 IFC array was primed and loaded onto a HX machine. Samples were then prepared with TaqMan Fast Universal PCR Master Mix (Life Technologies PN435042) and loaded on one end of the array. 20X TaqMan^®^ Gene Expression assays were mixed with 2X Assay Loading Reagent (Fluidigm PN 100-7611) and were loaded on the other end of the array. The 96.96 IFC was loaded onto the HX machine and finally transferred to Biomark HD machine. Results were analyzed using Excel and Qlucore for further analysis.

### Sirius Red, H&E and Masson’s Trichrome staining

Samples embedded in paraffin were mounted onto glass slides (VWR Micro Slides Catalog no 48311-703) using a rotary microtome (Sakura^®^ Accu-Cut SRM) at a thickness of 5 µm. After drying slides overnight at 37°C, slides were stored at RT until further staining with Sirius Red. Prior to using the Picro-Sirius Red Stain Kit (Abcam Catalog No. ab150681), samples were deparaffinized. Briefly, slides were dipped in two different Xylene substitute (Fisherbrand Safe Clear II Catalog No.044-192) consecutively for 5 and 10 min. Then slides were transferred into two washes with 100% ethanol for 3 min each. This is followed by two washes with 95% ethanol for 3 min each and a final rehydration step using two washes with deionized water for 5 min each. Following dehydration, slides are stained with Picro-Sirius Red for one hour at RT. Then, the slides are rinsed in acetic acid wash twice for one minute each. Finally, slides are rinsed in 100% ethanol 3 times for 1 min each and mounted in resin. Slides were also stained with Massons Trichrome using Trichrome Stain (Abcam Catalog No. ab150686). Slides were deparaffinized and hydrated as described previously. The Bouins protocol was not followed. Instead, slides were washed in Weigert’s Hematoxylin stain for 5 min at RT followed by a wash of deionized water. Then slides were washed in Biebrich Scarlet/Acid Fuchsin solution for 15 min then rinsed in deionized water. Samples were then washed in Phosphomolybdic/ Phosphotungstic Acid Solution for 10-15 min and placed in Aniline Blue solution for another 10-15 min. Samples were rinsed in distilled water, 1% acetic acid solution for 3-5 min, 95% ethanol twice for 30 sec each, and then 100% ethanol twice for 30 sec each. A final wash using Xylene substitute followed prior to final mounting with synthetic resin.

Samples for H&E staining were first deparaffinized and hydrated. Slides were placed in Hematoxylin for 3 min; washed in deionized water twice for 5 min; placed in SelecTech^®^ Define MX-aq (Leica Catalog No. 3803598) for 3 min; washed in water for 1 min; washed in 70% ethanol for 30 sec; stained with eosin for 30 sec; washed in 100% ethanol thrice for 1 min each and placed in Xylene Substitute twice for 3 min each. Slides were then covered slipped.

### Immunohistochemistry

Frozen sections in OCT were obtained using a Cryostat Thermo Scientific Microm HM550. Sections were mounted onto slides (VWR Micro Slides Superfrost Plus Catalog No 48311-703) at a thickness of 5 µm. Slides were stored at -20°C until further staining. For anti-mouse collagen staining, individual slides were thawed and dried for 20 min at RT; fixed using acetone at -20°C for 20 min; dried again at RT for 20 min; washed in TBS for 5 min; blocked for two hours at RT with Donkey anti-mouse IgG (Jackson Labs Catalog No.715-007-003); washed 3 times with TBS-T (.05% Tween) for 2 min each; stained and incubated in a humidified chamber with rabbit anti-mouse Collagen I overnight at 4°C (1:100 dilution Abcam Catalog No. ab34710); washed 3 times in TBS-Tween (.05% Tween) for 5 min each; stained with secondary antibody donkey anti-rabbit AF647 (Abcam Catalog No. 150075) for 1 hour at RT; rinsed 3 times with TBS-Tween for 5 min each; and finally mounted using Fluoroshield with DAPI (Sigma Catalog No. F6057-20ml). In conjunction with collagen I staining, anti mouse GFAP staining (astrocyte marker) was used. Anti-mouse GFAP AF594 (1:100 dilution Biolegend Catalog No. 644708).

## Notes

### Competing Interest Statement

The authors have declared no competing interest.

